# Easy-Multiome enables joint profiling of gene expression and chromatin accessibility in single cells

**DOI:** 10.64898/2026.07.15.738754

**Authors:** Xuan Zhang, Mark A.A. Minow, Robert J. Schmitz

## Abstract

Gene expression and chromatin accessibility provide complementary insights into the regulatory mechanisms that define cell states. Although methods for jointly profiling these modalities exist, plant applications remain limited because of complex workflows, inconsistent performance, and prohibitive costs. Here, we present easy-Multiome, a streamlined single-cell multiomic workflow that integrates a single *in situ* reverse transcription step into the standard droplet-based scATAC-seq protocol. Using easy-Multiome, we profiled more than 20,000 nuclei from maize seedlings generating paired gene expression and chromatin accessibility data, with approximately 90% of nuclei containing both high-quality RNA and chromatin accessibility profiles. The resulting transcriptome data resolved 16 clusters corresponding to nine major maize seedling cell types and enabled direct characterization of cell-type-specific chromatin accessibility from the same nuclei. Furthermore, easy-Multiome simultaneously captured cell-type-specific transcription factor expression and the accessibility of their cognate DNA-binding motifs, providing direct links between transcriptional programs and regulatory landscapes. Together, these results demonstrate that easy-Multiome enables robust and efficient joint profiling of plant gene expression and chromatin accessibility while requiring only minimal modifications to existing droplet-based scATAC-seq workflows.

## Introduction

Recent advances in single-cell sequencing technologies have enabled high-resolution characterization of diverse cellular modalities, including gene expression^1,2^, chromatin accessibility^3-5^, and protein abundance^6^. These technologies have provided unprecedented insights into developmental processes, tissue organization, and responses to environmental stimuli across diverse organisms^7-9^. Single-cell or single-nucleus RNA sequencing (scRNA-seq and snRNA-seq) have become the predominant method for characterizing cell identities and transcriptional states, particularly in plants^10,11^. Nevertheless, transcriptional profiles represent only a portion of gene regulation and the regulatory mechanisms that determine cell states and responses.

Chromatin accessibility provides complementary information by identifying potential *cis*-regulatory elements, containing transcription factor binding sites, that govern gene expression^12^. Consequently, single-cell ATAC-seq (scATAC-seq) has become a powerful approach for investigating gene regulatory mechanisms at the cellular level^13^. Plant chromatin accessibility profiling has discovered cell-type-specific regulatory elements, inferring transcription factor activity, and identifying regulatory programs behind development, environmental adaptation, and agronomic traits^3,14-16^. Integrating chromatin accessibility with gene expression provides a more comprehensive understanding of cellular gene regulation by linking *cis*-regulatory elements with downstream transcription. Such integrative analyses have reconstructed cell-type-specific gene regulatory networks, identified key transcriptional regulators, and characterized regulatory mechanisms^7,8^. However, chromatin accessibility and gene expression are often measured in separate experiments, requiring computational integration to associate the two modalities. These approaches rely on statistical matching of cells, batch effect corrections and may introduce uncertainty or bias, particularly when one dataset contains rare cell types or cellular states that are absent or underrepresented in the other^17,18^.

To overcome these limitations, single-cell multiomic technologies have been developed to simultaneously profile multiple molecular modalities from the same cell^19-21^. Chromatin accessibility and gene expression multiomics, directly link between *cis*-regulatory elements, and putative transcription factor activity, to cellular transcription, substantially improving the dissection of gene regulatory mechanisms. Among these methods, ISSAAC-seq (*in situ* SHERRY After ATAC-seq) demonstrated that chromatin accessibility and gene expression can be jointly profiled using droplet-based scATAC-seq platforms^21^. However, existing multiomic approaches involve multiple enzymatic reactions, repeated nuclei manipulation, and complex library construction procedures, which increase experimental complexity and reduce robustness. These challenges are particularly pronounced in plants, where nuclei isolation is often tissue dependent and very susceptible to damage during repeated processing. Consequently, current methods have seen relatively limited application across diverse plant species and tissues.

Here, we present easy-Multiome, a streamlined strategy for simultaneous profiling of chromatin accessibility and gene expression in single nuclei. By introducing a single *in situ* reverse transcription step into the standard droplet-based scATAC-seq workflow, easy-Multiome generates paired chromatin accessibility and transcriptome data. These minimal modifications to existing protocols retain library complexity while minimizing damage to fragile nuclei. Using more than 20,000 nuclei from maize seedlings, we demonstrate that easy-Multiome produces high-quality chromatin accessibility and gene expression profiles comparable to normal single-modality methods. We further show that the paired measurements accurately resolve major cell types and directly link cell-type-specific gene expression with their corresponding chromatin accessibility landscapes, providing a simple and broadly applicable approach for single-cell multiomic analysis.

## Results

### Easy-Multiome workflow overview

In the standard droplet-based scATAC-seq workflow from 10X Genomics^22,23^ **(Fig. 1A)**, isolated nuclei are directly tagmented by a Tn5 transposase loaded with standard Nextera adapters. The tagmented nuclei are subsequently loaded into a microfluidic device, where individual nuclei are encapsulated into droplet along with one barcoded gel beads. This process enables the profiling of single-cell accessible chromatin regions. In the easy-Multiome workflow **(Fig. 1B, Supplementary Methods)**, nuclei first undergo reverse transcription using primers containing a partial TruSeq adapter, a 10-bp unique molecular identifier (UMI), and a poly-T sequence. The nuclei are then subjected to Tn5 tagmentation, which simultaneously tagments accessible chromatin and RNA-DNA hybrids. Importantly, both reverse transcription and tagmentation are performed entirely *in situ* within intact nuclei, eliminating the need to physically separate chromatin and RNA.

**Fig. 1.**
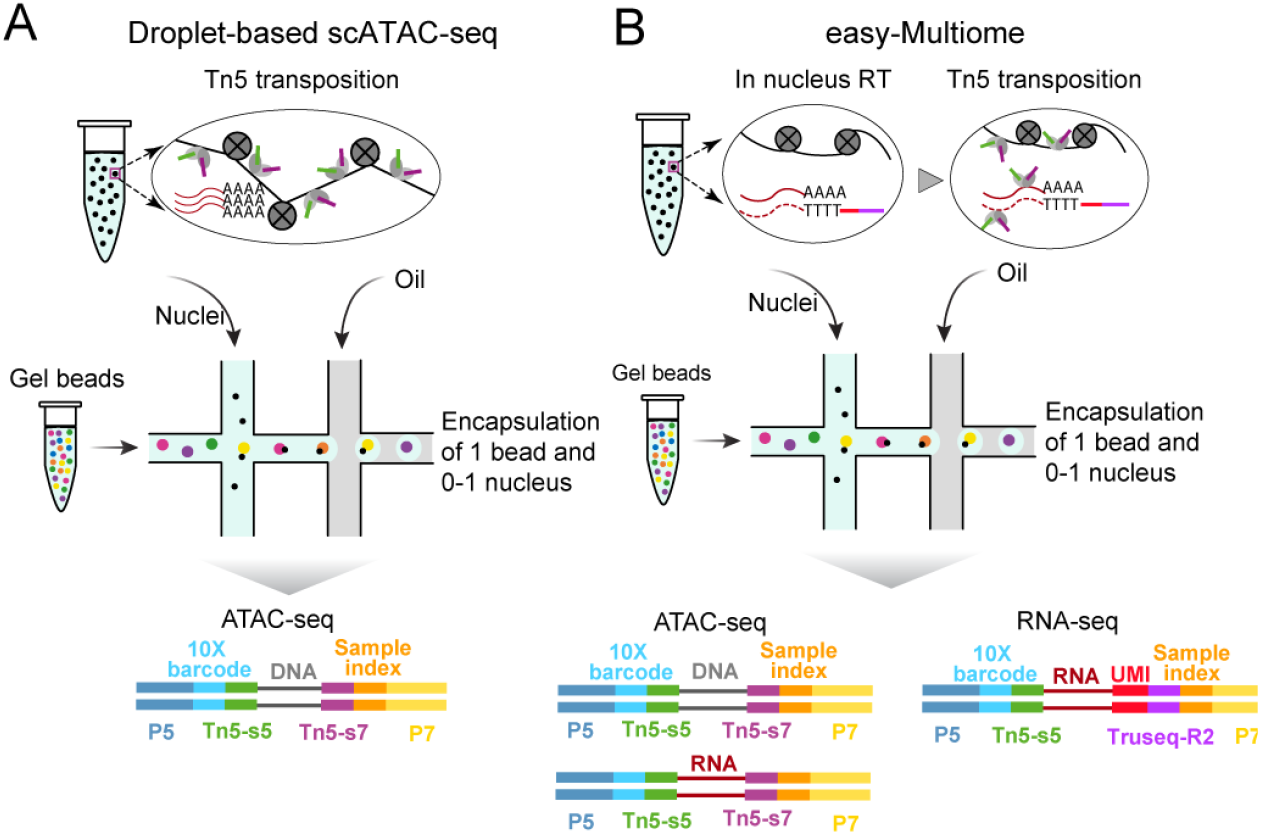
Easy-Multiome profiles chromatin accessibility and gene expression within the same nucleus. **(A)** Schematic overview of the standard droplet-based 10X Genomics single-cell ATAC-seq (scATAC-seq) workflow, in which nuclei are tagmented by Tn5 prior to droplet-based barcoding. **(B)** Schematic overview of the easy-Multiome workflow. Following nuclei isolation, an in situ reverse transcription step is introduced before Tn5 tagmentation, enabling simultaneous capture of gene expression and chromatin accessibility information from the same nucleus within the standard droplet-based scATAC-seq framework.

Following tagmentation, the workflow converges with the standard droplet-based scATAC-seq protocol, in which nuclei are encapsulated into droplets for single-cell barcoding. However, after recovery, and preamplification and the amplified library is divided equally into two fractions. One fraction is amplified with Nextera primers to generate the chromatin accessibility (ATAC-seq) library, whereas the other fraction is amplified using a combination of Nextera and TruSeq primers to generate the gene expression (RNA-seq) library. It is worth noting that RNA fragments with Nextera adapters on both ends can occasionally appear in the ATAC-seq data. However, these fragments predominantly originate from exonic regions and are therefore readily distinguished from ATAC-seq fragments, which are typically enriched in regulatory elements located in intronic and intergenic regions. Consequently, RNA-derived fragments can be efficiently removed by avoid the peaks overlap with exonic regions during downstream analysis by excluding peaks that overlap exonic regions, with minimal impact on chromatin accessibility profiling.

### Profiling gene expression and chromatin accessibility in the same cell using easy-Multiome

To evaluate the performance of easy-Multiome, we profiled maize seedlings with two biological replicates. We first examined the RNA and ATAC data as pseudo-bulk tracks using a genome browser **(Fig. 2A)**. The ATAC signal generated by easy-Multiome closely resembled that of scATAC-seq from 10X Genomics^3^, showing comparable Tn5 integration coverage across intergenic regulatory regions. The exception being a modest increase in gene-body tagmentation, caused by the small proportion of doubly Nextera tagged RNA-derived fragments. The RNA signal was predominantly within gene bodies, with a 3’ bias, consistent with snRNA-seq data from 10X Genomics^3^. Together, these results indicate that easy-Multiome generates high-quality chromatin accessibility and gene expression profiles.

**Fig. 2.**
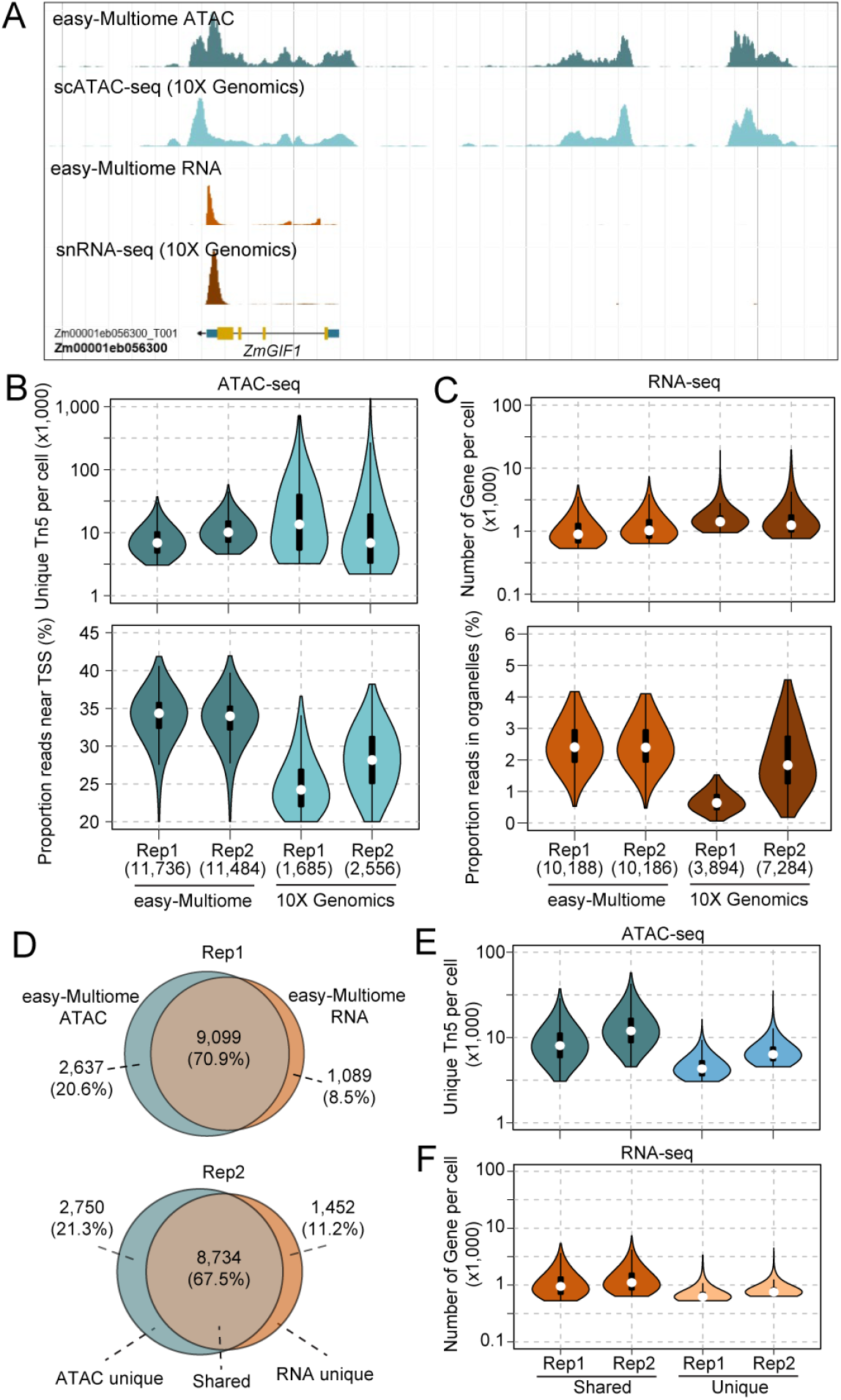
Quality assessment of easy-Multiome. **(A)** Pseudobulk genome browser tracks showing ATAC and RNA read coverage across the maize *ZmGIF1* locus generated by easy-Multiome. scATAC-seq from 10X Genomics and snRNA-seq datasets from the same tissue are shown for comparison. **(B)** Distribution of unique Tn5 insertions per nucleus and transcription start site (TSS) enrichment for nuclei profiled by easy-Multiome and conventional 10X Genomics scATAC-seq. **(C)** Distribution of the number of genes detected per nucleus and the proportion of organelle-derived transcripts for nuclei profiled by easy-Multiome and conventional 10X Genomics snRNA-seq. **(D)** Venn diagrams showing the overlap between high-quality ATAC and RNA nuclei identified by shared cell barcodes in the two biological replicates. **(E)** Distribution of unique Tn5 insertions for nuclei with both high-quality RNA and ATAC data (Shared) and nuclei with only high-quality ATAC data (Unique). **(F)** Distribution of the number of genes detected per nucleus for nuclei with both high-quality RNA and ATAC data (Shared) and nuclei with only high-quality RNA data (Unique).

We next evaluated data quality at the single-cell level for the ATAC and RNA data independently. Specifically, For the ATAC data, we assessed the number of unique Tn5 insertions per nucleus and transcription start site (TSS) enrichment **(Fig. 2B)**. For the RNA data, we evaluated the number of genes detected per nucleus and the proportion of reads mapping to organellar genes **(Fig. 2C)**. After quality control, we retained 11,484-11,736 ATAC nuclei and 10,168-10,188 RNA nuclei. Overall, the data quality was comparable to that of 10X Genomics scATAC-seq and snRNA-seq datasets generated from maize seedlings^3^.

To determine how efficiently easy-Multiome captures both molecular modalities within the same nucleus, we matched high-quality RNA and ATAC profiles using shared cell barcodes. Approximately 90% of high-quality RNA nuclei were paired with high-quality ATAC profiles in both biological replicates **(Fig. 2D)**. For example, among the 10,188 high-quality RNA nuclei in the first replicate, 9,099 (89.3%) also passed ATAC quality control. Notably, nuclei with both high-quality RNA and ATAC data generally exhibited higher data quality than nuclei with only a single modality, as reflected by greater numbers of unique Tn5 insertions and detected genes **(Fig. 2E,F)**. Together, these results demonstrate that easy-Multiome efficiently profiles gene expression and chromatin accessibility simultaneously in the same nucleus.

### Characterization of cell type-specific gene expression and chromatin accessibility

Because most cell type annotation is based on the expression pattern of cell-type-specific marker genes, which provides a more direct representation of cell state and identity than chromatin accessibility, we used the RNA data as the primary reference for cell-type annotation. We first integrated the snRNA-seq data from two biological replicates using the *Seurat* workflow and identified 16 distinct clusters **(Fig. 3A)**. Cell identities were assigned based on a curated list of published maize seedling marker genes^24^. Manual examination of marker gene expression further grouped these clusters into nine major cell types: epidermis, mesophyll, bundle sheath, pith parenchyma, cortex, vascular parenchyma, phloem, xylem and procambium **(Fig. 3B)**.

**Fig. 3.**
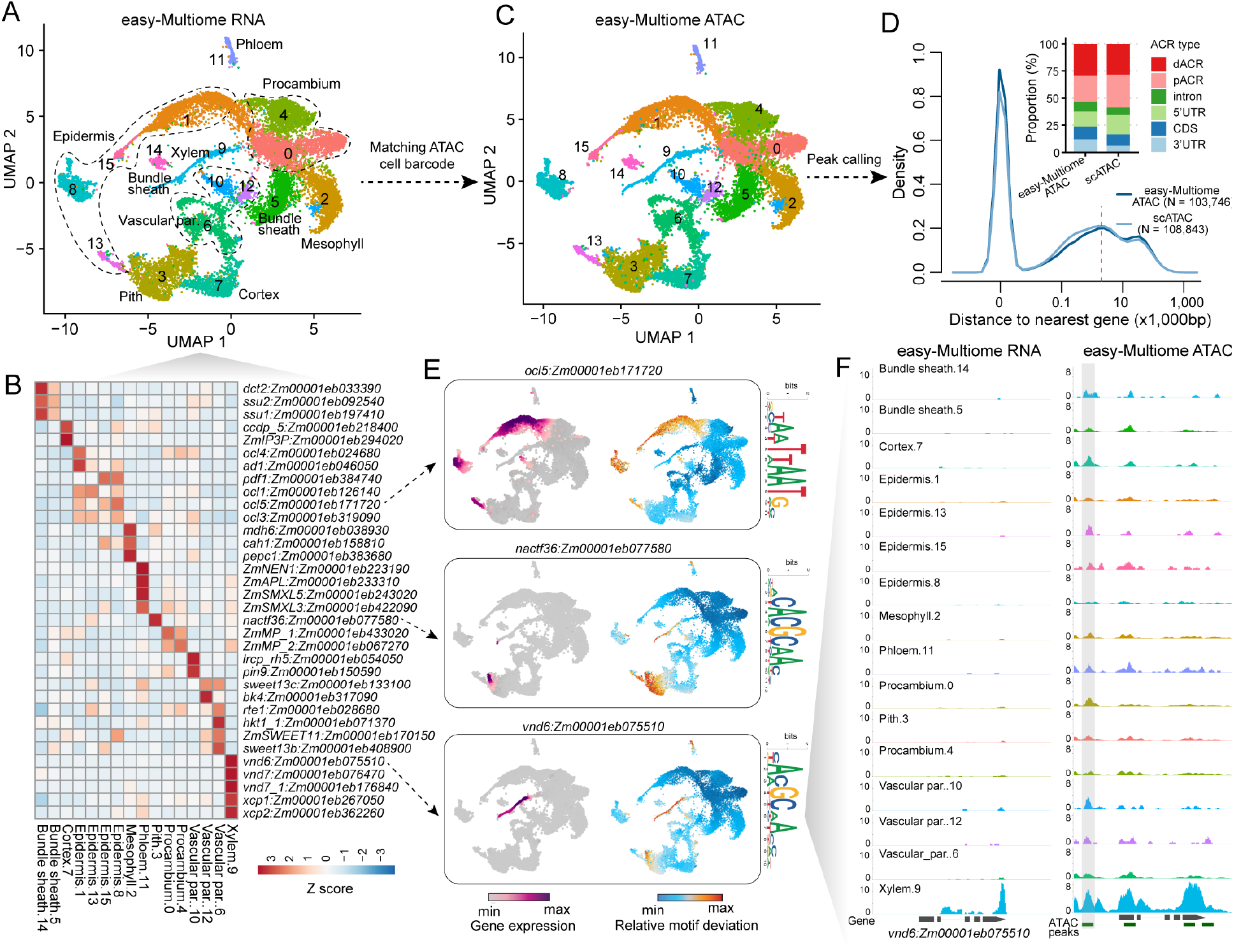
Characterization of cell type-specific gene expression and regulatory landscapes using easy-Multiome. **(A)** UMAP visualization of RNA profiles from all nuclei, colored by transcriptional clusters and annotated with predicted cell types. **(B)** Relative expression of representative marker genes across the identified cell types. **(C)** UMAP visualization of ATAC profiles from nuclei matched to RNA profiles using shared cell barcodes, colored and annotated according to corresponding cell types. **(D)** Comparison of ACRs identified by easy-Multiome and scATAC-seq from 10X Genomics. Left: density distribution of the distance between ACRs and the nearest gene, with the 2-kb threshold indicated by a red dashed line. Upper-right: genomic distribution of ACRs classified into six categories based on their location relative to gene annotations. **(E)** UMAP feature plots showing cell type-specific RNA expression (left) and corresponding transcription factor motif deviation scores derived from ATAC data (right) for representative transcription factors s, including OCL5, NACTF36, and VND6. **(F)** Pseudobulk RNA expression (left) and Tn5 integration signal (right) surrounding the xylem marker gene VND6, demonstrating the correspondence between cell type-specific gene expression and chromatin accessibility.

To characterize chromatin accessibility across cell types, we assigned ATAC profiles to the annotated cell types by matching shared cell barcodes between the RNA and ATAC datasets **(Fig. 3C)**. Across all cell types, we identified 103,746 accessible chromatin regions (ACRs). Because easy-Multiome ATAC libraries may contain certain proportion of RNA-derived fragments, we first evaluated the quality of the identified ACRs by comparing their genomic distribution with a previously published maize seedling scATAC-seq atlas^24^. The distance of ACRs to the nearest gene showed a distribution highly consistent with the published dataset **(Fig. 3D)**, indicating that easy-Multiome reliably identifies accessible regulatory regions in maize seedling.

We next classified ACRs according to the location of their summit relative to gene annotations into six categories: 5′ untranslated region (5′ UTR), coding sequence (CDS), 3′ untranslated region (3′ UTR), intron, proximal ACRs (pACRs; within 2 kb of a gene), and distal ACRs (dACRs; located more than 2 kb from the nearest gene). As expected, easy-Multiome identified a higher proportion of ACRs overlapping gene bodies than scATAC-seq from 10X Genomics, primarily because of increased 3′ UTR signals (12.10% versus 6.02%) derived from the RNA-seq signal. In contrast, the proportion of distal ACRs remained highly similar between the two datasets (29.28% versus 28.42%), demonstrating that easy-Multiome retains robust detection of distal regulatory regions. Because functional *cis*-regulatory elements are predominantly located within intergenic regions, promoters (overlapping 5’UTR), and introns^25^, ACRs overlapping CDS and 3′ UTR regions were excluded from subsequent analyses to minimize potential RNA-derived artifacts.

Transcription factors establish cell identity by binding to specific *cis*-regulatory elements and controlling cell type-specific gene expression^26^. Therefore, we expected the expression of cell-type-specific transcription factors to correlate with increased chromatin accessibility of their corresponding DNA-binding motifs. Consistent with this expectation, easy-Multiome simultaneously captured both transcription factors expression and motif chromatin accessibility within the same cell types **(Fig. 3E)**. For example, *OCL5* (*Outer Cell Layer 5*; *Zm00001eb171720*)^27^, a member of the HD-ZIP IV transcription factor family that regulates epidermal cell specification and cuticular wax biosynthesis, was specifically expressed in epidermal cells, where its binding motif also displayed elevated accessibility. Likewise, *VND6* (*VASCULAR-RELATED NAC-DOMAIN 6*; *Zm00001eb075510*)^28^, a key regulator of xylem differentiation, showed developing xylem-specific expression accompanied by increased accessibility of its cognate motif in xylem cells. Easy-Multiome additionally revealed xylem-specific ACRs surrounding the *VND6* locus, highlighting candidate *cis*-regulatory regions that may contribute to *VND6* cell type-specific expression **(Fig. 3F)**.

Together, these results demonstrate that the addition of a single in-nucleus reverse transcription step to the standard scATAC-seq workflow enables easy-Multiome to simultaneously generate high-quality gene expression and chromatin accessibility profiles from the same nucleus while preserving accurate identification of cell types and their regulatory landscapes.

## Discussion

Simplicity is an important but often underappreciated aspect of method development. For a technology to be broadly adopted, it must be both robust, easy to implement and cost effective. ISSAAC-seq is an alternative approach for simultaneous profiling of gene expression and chromatin accessibility using droplet-based scATAC-seq platforms^21^, including the 10X Genomics workflow. However, its protocol involves multiple sequential nuclei treatments, including chromatin tagmentation, *in situ* reverse transcription, RNA-DNA hybrid tagmentation, exonuclease I digestion, and gap filling before cell barcoding. These procedures require four separate enzymatic reactions interspersed with multiple centrifugation and wash steps, which may compromise nuclei quality and reduce nuclei recovery. Consequently, the workflow may be more dependent on highly intact nuclei, such as those isolated from cultured cell lines, potentially limiting its application to fragile or difficult-to-isolate tissues (https://xichenlab.com/posts/2022-09-21-issaac/). Consistent with this limitation, our numerous attempts to apply ISSAAC-seq to maize seedlings were unsuccessful, as the resulting RNA libraries lacked sufficient complexity for reliable cell-type identification. In contrast, easy-Multiome requires only a single additional *in situ* reverse transcription step comparing the standard scATAC-seq workflow. Performing reverse transcription immediately after nuclei isolation helps preserve RNA integrity before prolonged incubations, while minimizing nuclei manipulation and wash steps, improving method robustness.

A key advantage of easy-Multiome is its flexibility. Because easy-Multiome modifies only the nuclei preparation step while retaining the standard droplet-based scATAC-seq workflow, it can be readily integrated with fluorescence-activated cell sorting and multiple droplet-based platforms that use Nextera capture chemistry, including 10x Genomics, Bio-Rad, and HyDrop. Beyond maize, we have successfully applied easy-Multiome to several additional plant species, including *Medicago truncatula*, common bean (*Phaseolus vulgaris*), soybean (*Glycine max*) and switchgrass (*Panicum virgatum*), consistently obtaining high-quality RNA-seq data.

Despite these advantages, easy-Multiome has several limitations. Because reverse-transcribed RNA molecules can occasionally be incorporated into the ATAC library, RNA-derived fragments may be misidentified as ACRs, particularly in species with incomplete genome annotations. Although these fragments can usually be distinguished by their enrichment within gene bodies and removed during downstream analysis, excessive RNA carryover may reduce ATAC-seq data quality. For example, in soybean hypocotyl we observed substantially higher gene-body coverage than intergenic coverage, suggesting extensive RNA-derived signal that could interfere with accurate ACR identification. We therefore recommend evaluating libraries at the bulk level before scaling the protocol to new species or tissue types and optimizing experimental conditions when excessive RNA-derived signal is observed.

The simplicity of the easy-Multiome workflow also suggests opportunities for extending this strategy to other single-cell sequencing applications. For example, easy-Multiome should be compatible with approaches such as GoT-ChA (Genotyping of Targeted Loci with Chromatin Accessibility)^29^, enabling the simultaneous measurement of genotype, gene expression, and chromatin accessibility within the same cell. Furthermore, its streamlined design may provide a foundation for developing ultra-scalable single-cell multiomic approaches, similar to SUM-seq^30^, capable of profiling hundreds of samples at the million-cell scale.

Here, we present easy-Multiome, a simple modification of the standard droplet-based scATAC-seq workflow that enables paired multiomic profiling with minimal changes to existing protocols. By integrating reverse transcription directly into the existing workflow, easy-Multiome simultaneously measures chromatin accessibility and gene expression from the same nucleus without requiring specialized instrumentation or extensive protocol modifications. Its streamlined design lowers the technical barrier to single-cell multiomic profiling while maintaining compatibility with established scATAC-seq workflows. We envision that easy-Multiome will make paired multiomic analysis more broadly accessible to the plant research community and serve as a versatile platform for future innovations in scalable, multimodal single-cell genomics.

## Methods

### Growth conditions and tissue collection

The maize seed for genotype B104 were sown in Sungro Horticulture professional growing mix (Sungro Horticulture Canada Ltd.). Soil was saturated with tapwater and placed under a 50/50 mixture of 4100K (Sylvania SupersaverCoolWhite Delux F34CWX/SS, 34W) and 3000K (GE Ecolux w/ starcoat, F40CX30ECO, 40W) lighting. Seedlings were grown under a photoperiod of 16 hours of light, eight hours of dark. The temperature was approximately 25℃ during light hours with a relative humidity of approximately 54%. Seedling tissue extending 1 cm on either side of the first visible node (2 cm total) were harvested between 8 and 9 AM six days after sowing. Around 3-5 seedling were used for each replicate.

### easy-Multiome protocol

A detailed step-by-step easy-Multiome protocol, including the required reagents, equipment, and oligonucleotide sequences, is provided in the Supplementary Methods and Supplementary Table S1. Briefly, Nuclei were isolated from fresh plant tissues by mechanical chopping in nuclei isolation buffer, followed by filtration, Percoll gradient purification, and washing. Approximately 20,000 purified nuclei were subjected to in situ reverse transcription using a poly(dT)-containing reverse transcription primer and Maxima H Minus Reverse Transcriptase. Following reverse transcription, nuclei were washed to remove excess RNA and reaction components before proceeding with the standard 10X Genomics Chromium Single Cell ATAC library preparation workflow. Tagmentation, GEM generation, linear amplification, and DNA purification were performed according to the manufacturer’s protocol (10x Genomics Chromium Next GEM Single Cell ATAC Reagent Kit). RNA- and ATAC-derived libraries were independently amplified using library-specific index primers, with the number of PCR cycles determined by qPCR to minimize amplification bias. Final libraries were purified using SPRI beads and sequenced on an Illumina platform.

### Library preparation and sequencing

easy-Multiome libraries were prepared using the Chromium scATAC v2 (Next GEM) kit from 10X Genomics, following the manufacturer’s instructions. (10X Genomics, CG000496_Chromium_NextGEM_SingleCell_ATAC_ReagentKits_v2_UserGuide_RevB). The leftover nuclei after loading to ChIP-H were diluted, stained with DAPI, and nuclei quality and density was evaluated with a hemocytometer under a microscope. The final libraries were sequenced using an Illumina NovaSeq X Plus in the dual-index mode.

### RNA sequencing data analysis

For RNA-seq data, Index 2 and the first 10 bp of Read 2 were concatenated to generate a new FastQ file containing the cell barcode and UMI (CB_UMI.fastsq). Then the CB_UMI.fastq and Read 1 files were used to generate the gene expression matrix by mapping the reads to B73 AGP v5 maize reference genome^31^ with STARsolo^32^. We specified the following parameters in STARSolo to filter the UMI, filter empty cells, and count unique mapping reads: --soloUMIfiltering MultiGeneUMI_CR, --soloCellFilter EmptyDrops_CR. The filtered expression data was analyzed using the Seurat (v4) R package^33^. Potential low-quality nuclei or empty droplets were filtered. Specifically, cells with gene counts below a threshold calculated as the median gene count minus two times the median absolute deviation, and cells with UMI counts less than the lower 10% percentile of total UMI counts, were filtered out. Additionally, cells with organelle gene counts comprising more than 15% of the total gene count were excluded. The preprocessed datasets were normalized using SCTransform before the RunPCA for principal component analysis (PCA). Subsequently, the doublets were identified by the DoubletFinder R package^34^, and removed from downstream analysis. We prepared two replicates for each library and integrated them using the Harmony R package. The integrated dataset was then processed using RunUMAP (reduction = “harmony”, dims = 1:20) for Uniform Manifold Approximation and Projection (UMAP) dimension reduction, FindNeighbors (reduction = “harmony”, dims = 1:30) to obtain the Nearest-neighbor graph, and FindClusters to identify distinct cell populations. The cluster 0 from initial clustering were excluded for downstream analysis because exhibited a low number of detected genes and UMIs, showed broad correlation with the expression profiles of other clusters, and lacked clear cell type specificity.

### ATAC sequencing data analysis

The raw data processing followed the previously described method^3^. In brief, raw BCL files were demultiplexed and converted into fastq format using the default settings of the 10X Genomics tool cellranger-atac makefastq (v1.2.0). Initial read processing, including adaptor/quality trimming, mapping, and barcode attachment/correction, was carried out with cellranger-atac count (v1.2.0) using the B73 AGP v5 maize reference genome^31^. Properly paired mapped reads with a mapping quality greater than 30 were retained using samtools view (v1.6; -f 3 -q 30)^35^, while also retaining reads with alternate hit XA tags to avoid biasing downstream analysis due to the whole genome duplication events during soybean evolution. Duplicate fragments were collapsed on a per-nucleus basis using picardtools (http://broadinstitute.github.io/picard/) MarkDuplicates (v2.16; BARCODE_TAG=CB REMOVE_DUPLICATES=TRUE). Reads mapping to mitochondrial and chloroplast genomes were counted for each barcode and then excluded from downstream analysis. BAM alignments were then converted to single base-pair Tn5 integration sites in BED format by adjusting coordinates of reads mapping to positive and negative strands by +4 and -5, respectively, and retaining only distinct Tn5 integration sites for each individual barcode.

The R package *Socrates* was used for nuclei identification and quality control^3^. The BED file containing single base-pair Tn5 integration sites was imported into *Socrates* along with the maize GFF gene annotation and the genome index file. To identify bulk-scale ACRs in *Socrates*, the callACRs function was employed with the following parameters: genome size=8.0e8, shift=-75, extsize=150, and FDR=0.1. This step allowed us to estimate the fraction of Tn5 integration sites located within ACRs for each nucleus. Metadata for each nucleus were collected using the buildMetaData function, with a TSS (Transcription Start Site) window size of 1 kb (tss.window=1000). Sparse matrices were then generated with the generateMatrix function, using a window size of 500. High-quality nuclei were identified based on the following criteria: a minimum of 1,000 Tn5 insertion sites per nucleus, at least 20% of Tn5 insertions within 2 kb of TSSs, and at least 20% of Tn5 insertions within ACRs across all datasets. Additionally, a maximum of 20% of Tn5 insertions in organelle genomes was allowed.

### ACR identification

Following cell clustering and annotation with the RNA-seq data, peaks were identified using all Tn5 integration sites for each cluster by running MACS2 with non-default parameters: --extsize 150 --shift -75 -- nomodel --keep-dup all^36^. To account for potential bias introduced by read depth, we adjusted the q-value cutoffs based on the total Tn5 integration number in each cell type as follows: for less than 10 million integrations, we used --qvalue 0.1; for 10-25 million, we used 0.05; for 25-50 million, we used 0.025; for 50-100 million, we used 0.01; and for more than 100 million, we used 0.001. Peaks were then redefined as 500-bp windows centered on the peak coverage summit. To consolidate information across all clusters, we concatenated all peaks into a unified master list using a custom script^3^. The peak chromatin accessibility score was calculated based on the Tn5 integration count within the peak and then normalized using the cpm function in edgeR^37^. ACRs with less than 4 CPM in all cell types and its summit overlap with 3’UTR or CDS regions were removed from downstream analysis.

### Transcription factors Motif deviations score calculation

Transcription factors motif deviation scores of specific TF motifs among nuclei were estimated using chromVAR^38^ with the motif identified with DAP-seq in maize in previous study^39^. The input matrix for chromVAR was filtered to retain ACRs with a minimum of 10 fragments and cells with at least 100 accessible ACRs. We applied smoothing to the bias-corrected motif deviations for each nucleus, integrating them into UMAP embedding for visualization^3^.

## Acknowledgments

We would like to thank Yinxin Dong and Yangyang Xu for their assistance in maintaining an organized laboratory environment. This study was supported in part by resources and technical expertise from the Georgia Advanced Computing Resource Center, a partnership between the University of Georgia’s Office of Research.

## Author contributions

R.J.S., and X.Z. designed the research. X.Z. performed the experiments. X.Z., M.A.A.M. and R.J.S. analyzed the data. X.Z., M.A.A.M. and R.J.S. wrote the manuscript. All authors read and approved the final manuscript.

## Data availability

All datasets generated in this study are available at GEO (Accession number: GSE338572). All original code has been deposited at Github (https://github.com/schmitzlab/easy-Multiome). Any additional information in this paper is available from the lead contact upon request.

