## Supplementary method for "Easy-Multiome enables joint profiling of gene expression and chromatin accessibility in single cells"

### Supplementary Methods

**Easy-Multiome protocol.** The developed approach should be applicable easily with little modification from droplet-based single cell ATAC-seq (scATAC). We have also applied it to *Medicago truncatula*, common bean (*Phaseolus vulgaris*), soybean (*Glycine max*) and switchgrass (*Panicum virgatum*). Although we are consistently obtaining high-quality RNA-seq data, excessive RNA carryover may reduce ATAC-seq data quality, we therefore recommend evaluating libraries at the bulk level before scaling the protocol to new species or tissue types and optimizing experimental conditions when excessive RNA-derived signal is observed. A step-by-step protocol is presented below in detail.

I. **Adapter and primer design (Table S1):**
1) Reverse transcription (RT) oligo:

eRNA-Truseq-polyT (10 μM): CAGACGTGTGCTCTTCCGATCTNNNNNNNNNNTTTTTTTTTTTTTTTTTTTTTTTTTTTTTTVN

2) Index PCR/qPCR primer set (10 μM):

Partial-P5: 5'-AATGATACGGCGACCACCGAGA-3'

ATAC_N70X (Table S1):

5'-CAAGCAGAAGACGGCATACGAGATXXXXXXXXGTCTCGTGGGCTCGG-3'

RNA _N70X (Table S1):

5'-CAAGCAGAAGACGGCATACGAGATXXXXXXXXGTGACTGGAGTTCAGACGTGTGCTCTTCCGATCT-3'

**II. Nuclei isolation with quick purification**

1. Prepare 10 mL Nuclei Isolation Buffer Base (**NIB-base**) on ice, make fresh.

| Stock | Final | 10 mL |
| --- | --- | --- |
| 0.4M MES-KOH pH 5.4 | 10 mM | 250 uL |
| 2M KCl | 10 mM | 50 uL |
| 1M MgCl_2_ | 3 mM | 30 uL |
| **5M NaCl** | 10 mM | 20 uL |
| 2M sucrose | 0.25 M | 1.25 mL |
| H2O |  | to 10 mL |

2. Prepare **NIB-wash buffer** on ice (about 2 mL per sample):

| Stock | Final | 1 mL |
| --- | --- | --- |
| NIB-Base buffer |  | 1 mL |
| 10% BSA (Miltenyi Biotec 130-091-376) | 0.5% | 50 uL |
| Protector (Sigma 3335399001) (40U/uL) | 0.4U/uL | **5 uL** |
| SUPERase Inhibitor (ThermoFischer AM2696)(20 U/uL) | 0.2U/uL | **5 uL** |
| RiboLock (ThermoFisher EO0382)(40 U/uL) | 0.4U/uL | **5 uL** |

3. Prepare **NIB-cutting Buffers**: about 1 mL per sample:

| Stock | Final | 1 mL |
| --- | --- | --- |
| NIB-Base buffer |  | 1 mL |
| 1M Spermine | 0.1 mM | 0.2 uL |
| 1M Spermidine | 0.5 mM | 0.5 uL |
| 1M DTT | 1 mM | 1 uL |
| 10% 10% NP-40 | 0.5% | 50 uL |
| 10% PVP-40 | 0.5% | 50 uL |
| cOmplete™ Protease Inhibitor Cocktail EDTA free (Roche 11836170001; 50 X, 1 tablet in 200 uL H_2_O) | 1x | 20 uL |
| Protector (Sigma 3335399001) (40U/uL) | 0.4U/uL | 10 uL |
| SUPERase Inhibitor (ThermoFischer AM2696)(20 U/uL) | 0.2U/uL | 10 uL |
| RiboLock (ThermoFisher EO0382)(40 U/uL) | 0.4U/uL | 10 uL |
| 40% glyoxal | 3% | 75 uL |
| 100% acetic acid | 0.75% | 7.5 uL |

4. Prepare 1mg/mL DAPI (1,000 X): Dissolve 1mg DAPI (sigma Cat. D9542) in 1mL ddH2O, store at -20 ℃
5. Add 500 μL of **NIB-cutting** buffer (adjusted based on tissue amount) to the Petri dish. Finely chop the tissue on ice with a fresh razor blade for 1 minute, keeping the homogenized tissue concentrated in the center of the dish.
6. Filter the homogenate through a 40-μm cell strainer into a pre-chilled tube.
7. Incubate the nuclei suspension on ice for 10 minutes to complete nuclei lysis/fixation.
8. Pellet the nuclei by centrifugation at 500 × g for 5 minutes at 4°C. Carefully remove and discard the supernatant without disturbing the pellet.
9. Gently resuspend the nuclei pellet in 500 μL **NIB-wash buffer** (without NP-40) and filter the suspension through a 20-μm cell strainer.
10. Carefully layer the filtered nuclei suspension on top of 500 uL 35% Percoll solution (175 uL Percoll + 325 uL **NIB-wash buffer**) without disturbing the interface. Centrifuge at 500 × g for 8 minutes at 4°C. After centrifugation, most green debris will remain near the top of the Percoll layer, while the nuclei will pellet at the bottom of the tube.
11. Wash the nuclei once with 500 uL **NIB-wash buffer**.
12. Resuspend the nuclei pellet in 100 μL NIB wash buffer and mix gently until the suspension is homogeneous.
13. Dilute 5 μL of the nuclei suspension 1:10 with NIB wash buffer, stain with DAPI, and determine the nuclei concentration using a fluorescence microscope or cell counter.
14. Transfer 20,000 nuclei (X μL) into a new PCR tube and bring the volume to 200 μL with NIB wash buffer. Centrifuge at 500 × g for 3 minutes at 4°C. Carefully remove the supernatant, leaving approximately 5 μL in the tube without disturbing the nuclei pellet.

**III. *In situ* reverse transcription.**

15. Prepare a 25-μL RT reaction for each sample, including the 5 μL nuclei suspension. Mix gently by slowly pipetting up and down 10–20 times.

| Stock | Final | 25 uL |
| --- | --- | --- |
| 10 uM eRNA-Truseq-polyT | 2 uM | 5 uL |
| 5X NaCl RT buffer | 1x | 5 uL |
| Nuclease-free Water |  | 2.75 |
| 50% PEG8000 | 6% | 3 uL |
| 10mM dNTP(Thermo Scientific, R0192) | 0.5 mM | 1.25 uL |
| Maxima H minus Reverse  Transcriptase (Thermo Scientific, FEREP0753, 200 U/uL) | 10U/uL | 1.25 uL |
| RiboLock (ThermoFisher EO0382)(40 U/uL) | 0.4U/uL | 0.25 uL |
| Protector (Sigma 3335399001) (40U/uL) | 2U/uL | 1.25 uL |
| SUPERase Inhibitor (ThermoFischer AM2696)(20 U/uL) | 0.2U/uL | 0.25 uL |
| Nuclei | About 5uL | 5 uL |

16. Gently resuspend each nuclei pellet in 20 μL RT buffer by slowly pipetting up and down 20 times. Mix thoroughly while avoiding the introduction of bubbles.
17. Incubate the samples in a thermocycler at 25°C for 10 minutes, followed by 37°C for 15 minutes**.**

**IV. Easy-Multiome library preparation:**

18. Add 200 μL ice-cold 1× DNB buffer containing 0.5% BSA to each RT reaction and mix gently.
19. Centrifuge at 500 × g for 3 minutes at 4°C. Carefully remove and discard the supernatant.
20. Wash the nuclei by gently resuspending the pellet in 200 μL ice-cold 1× DNB buffer containing 0.5% BSA. Centrifuge at 500 × g for 3 minutes at 4°C.
21. Carefully remove the supernatant, leaving approximately 5 μL of nuclei suspension at the bottom of the tube.
22. Proceed with the standard 10X Genomics scATAC-seq protocol for tagmentation..
23. After 12 cycles of linear amplification in the GEMs, purify the DNA according to the 10x Genomics protocol. Elute the purified DNA in 2 × 22 μL of nuclease-free water.
24. Run qPCR to check the Cq for RNA and ATAC-seq lib.

- With the aim of determining PCR cycles prior to saturation occurs to mitigate amplification bias for the library construction, 1 μL of DNA sample from above was spared to perform the 10 μL real-time PCR (qPCR):

5 μL of 2x Luna® Universal qPCR Master Mix(NEB, cat# M3003)

0.25 μL Partial-P5

0.25 μL ATAC_N70X(ATAC) or RNA_N70X (RNA)

0.5 μL DNA Sample

4 μL Nuclease-Free Water

- Program Bio-Rad CFX96 qPCR system and perform the thermal profile as below:

Initial denaturation for 30s at 95°C

Denature for 15s at 95°C

30 cycles	Anneal primers for 30s at 60°C

Detect fluorescence	Extend DNA for 1 min at 72°C

4°C Hold

25. Perform library index amplification following the protocol described below.

**PCR reaction:**

| Component | volume |
| --- | --- |
| RNA_N70X/ATAC_N70X(10uM) | 2.5 uL |
| Partial P5 (10 uM) | 2.5 uL |
| Q5 High-Fidielity 2X Master Mix (NEB, M0492L) | 25 uL |
| DNA sample | 20 uL |

**PCR program for ATAC and RNA:**

|  | Temperature | Time | Number of cycles |
| --- | --- | --- | --- |
| Initial denaturation | 98C | 1 min | 1 |
| Denaturation | 98C | 20 sec | Cq |
| Annealing | 63C | 20 sec |  |
| Extension | 72C | 20 sec |  |
| Final extension | 72C | 1 min | 1 |
| Hold | 10C | hold | hold |

26. Perform the final cleanup using SPRI beads. Purify the ATAC library with 1.1× bead volume and the RNA library with 0.8× bead volume. Elute both libraries in 20 μL nuclease-free water.

### Supplementary Note:

In theory, there should be no sample or species limits for easy-Multiome, provided that high-quality nuclei can be isolated and work for scATAC-seq. We have successfully obtained high-quality nuclei for numerous plant species with the current nuclei isolation protocol, including maize, soybean, rice, wheat, barley, Arabidopsis, and poplar, across various tissue types such as leaf, root, nodule, hypocotyl, and developing seed. We recommend validating this nuclei isolation protocol by assessing nuclei quality under a microscope and evaluating the data quality through regular/non-single-cell easy-Multiome prepared from the isolated nuclei.
